# Pan-angiosperm analysis of the CLE signaling peptide gene family unveils paths, patterns, and predictions of paralog diversification

**DOI:** 10.1101/2025.06.04.657892

**Authors:** Iacopo Gentile, Miguel Santo Domingo, Sophia G Zebell, Blaine Fitzgerald, Zachary B. Lippman

## Abstract

The compositions of conserved gene families often vary widely between species, complicating predictions and experimental tests of shared versus distinct functions, especially in families shaped by extensive duplication, redundancy, and paralog diversification. The plant *CLV3/EMBRYO-SURROUNDING REGION* (*CLE*) small-signaling peptide family exemplifies these challenges. Although genetic studies in model systems have identified shared roles for a few *CLE* genes and species-specific redundancies, an evolutionary analysis of the entire family over deep time could empower predictive and experimental dissections of functions obscured by redundancy. We developed a scanning pipeline that *de novo* annotated *CLE* genes from 2,000 genomes representing 1,000 species, uncovering thousands of previously undetected family members and producing a comprehensive phylogenetic reconstruction and tracing of the family’s evolution and sequence diversification over 140 million years. Computational modeling of coding and cis-regulatory regions predicted lineage-specific asymmetries in paralog redundancy, stemming from ancestral amino acids in the functional core of the dodecapeptide and partial conservation of promoter elements. We tested these predictions using two genome-editing strategies in Solanaceae. Base-editing of deeply conserved residues in the CLV3 dodecapeptide and its paralogs across three species confirmed their critical roles in repressing stem-cell proliferation, and multiplex CRISPR knockouts of the 52 tomato *CLE* genes resolved pairwise and higher-order redundancies, revealing previously uncharacterized regulators of shoot architecture and plant size. These findings show how both peptide and cis-regulatory erosion shape *CLE* redundancy and provide a framework for detecting and translating deep evolutionary signals into testable genetic hypotheses across compositionally complex gene families.

## Introduction

High-throughput genome sequencing and computational genomics has transformed our understanding of gene family evolution across evolutionary timescales. Comparative analysis of genome composition has revealed dynamic and complex patterns of gene birth, death, and functional divergence. Gene families, formed and expanded through duplication events, exhibit remarkable variation in sequence, expression, and function across both distantly and closely related species (Masatoshi Nei 2013; Murat, Peer, and Salse 2012). The mechanisms driving this diversity operate through distinct evolutionary trajectories: initial redundancy following a duplication event typically degrades through mutational drift, often resulting in gene loss (pseudogenization). However, through mutational serendipity and under certain selective pressures, a duplicated gene (hereafter, paralog) may partition its functions with its ancestor (subfunctionalization), or acquire new roles (neofunctionalization) (Wagner 2008; Dittmar and Liberles 2011). Although these classical long-term endpoints of paralog functional evolution have been well documented across many lineages, the evolutionary trajectories and dynamics of paralogous gene diversification over shorter timescales are less understood (Lynch et al. 2001; Lynch and Conery 2003; Birchler and Yang 2022). Recent pan-genomic studies offer opportunities to capture a range of evolutionary timescales that can reveal how lineage-specific duplications diversify gene families in sequence and function (Benoit et al. 2025; W. He et al. 2025). In particular, deep evolutionary sampling through pan-genomics can reveal how lineage- and species-specific paralog redundancies arise and shift from the combined effects of coding and non-coding sequence variation (Kwon et al. 2022; Light and Kraulis 2004; Tvrdik and Capecchi 2006).

Following whole genome or local gene duplication events, redundancy among paralogs allows mutations to accumulate in coding and regulatory sequences, leading to unpredictable changes in initial redundancy relationships that can affect genotype-phenotype relationships. An integrated approach that combines comprehensive phylogenetic sampling using expansive pan-genomic data with predictive computational modeling via machine learning approaches has the potential to reveal the dynamics of how redundancy relationships evolve to shape gene families and their functional compositions. A remaining barrier, however, is incomplete and inconsistent gene annotations between reference genomes, which continues to obscure the full extent of genetic and allelic diversity, particularly in gene families that have undergone, and continue to undergo, frequent duplication and sequence evolution.

A striking example of these challenges is found in the plant *CLAVATA3/EMBRYO SURROUNDING REGION-RELATED* (*CLE*) small-signaling peptide gene family. *CLE* genes encode approximately 100 amino acid peptide sequences that are proteolytically processed into 12-amino-acid small signaling peptides (dodecapeptides). The dodecapeptides are glycosylated and then secreted to bind leucine-rich repeat receptor-like kinases (LRR-RLKs) on the surface of neighboring cells. These interactions mediate downstream signaling events that are critical for diverse developmental and physiological programs (Whitewoods 2021). However, high sequence divergence surrounding the functional dodecapeptides coupled with extensive copy number variation and challenges in detecting tissue- and cell-specific expression have confounded CLE family annotation and thus predictions that could permit systematic and comprehensive comparative functional analysis within and across species, especially paralog redundancy (Carbonnel, Falquet, and Hazak 2022; Carbonnel, Cornelis, and Hazak 2023). Indeed, a comprehensive mutational analysis of all *CLE* genes in the model *Arabidopsis thaliana* (hereafter Arabidopsis) revealed that most single-gene knockouts show no obvious phenotypes, suggesting widespread redundancy and compensatory relationships across family members that can only be revealed through high order genetics (Yamaguchi et al. 2017), as demonstrated in the meristem interactive signaling between tomato *SlCLV3* and *SlCLE9* (Rodriguez-Leal et al. 2019) and Arabidopsis *AtCLV3*, *AtCLE16*, and *AtCLE17* (Dao et al. 2022).

Here, we aggregated plant pan-genomic resources and developed a computational pipeline to rapidly identify and annotate *CLE* genes across 2,000 genomes representing 1,000 species and spanning 140 million years of evolution (De Bodt, Maere, and Van De Peer 2005). By integrating comparative phylogenetic analysis spanning ancient and recent evolutionary timescales, predictive computational modelling of the mutational landscape, and functional characterisation through CRISPR genome editing, we uncovered mechanisms underlying the maintenance and diversification of *CLE* paralog redundancy. Our findings demonstrate that resolving the long-term dynamics of coding and regulatory sequence evolution among gene family members can predict the architectures of complex paralog interactions, exposing redundancy relationships and enhancing the predictability of genome editing outcomes.

## Results and Discussion

### Pan-Angiosperm discovery and analysis of the CLE peptide

*CLE* genes encode precursor proteins typically less than 100 amino acids in length that consist of an N-terminal Golgi signal peptide, a variable domain with significant sequence divergence, and a C-terminal 12–amino acid CLE motif called a dodecapeptide (**Figure 1A**; Fletcher 2020). After proteolytic cleavage and release from the precursor, CLE dodecapeptides undergo post-translational modifications essential for activity (Whitewoods 2021; Jeong et al. 2024; Ohyama et al. 2009). By binding to and signaling through leucine-rich repeat (LRR) receptors, these small signaling peptides regulate numerous critical developmental and physiological programs, both conserved and species-specific (Fletcher 2020; Araya, Von Wirén, and Takahashi 2014). For instance, *CLV3* is a deeply conserved *CLE* family member that represses stem cell proliferation in the shoot apical meristem (Fletcher et al. 1999; Benoit et al. 2025; Rodriguez-Leal et al. 2019; Kwon et al. 2022), whereas the Arabidopsis *AtCLE14* and *AtCLE42* paralogous genes, regulate leaf senescence (Z. Zhang et al. 2022; Y. Zhang et al. 2022), and *MtCLE53* in the leguminous species *Medicago truncatula* controls root nodulation (Karlo et al. 2020) (**Figure 1A**).

**Figure 1.**
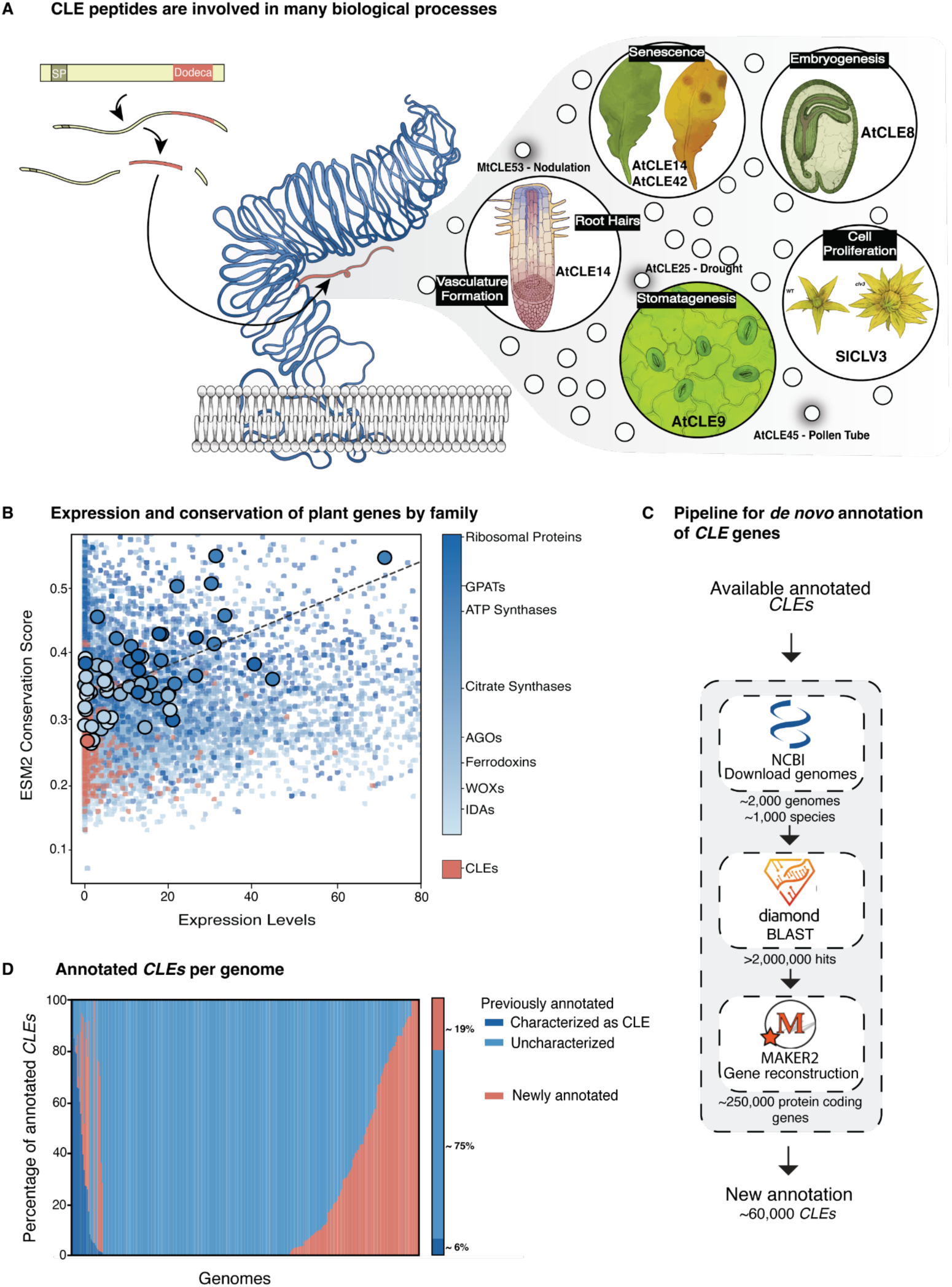
*De novo* annotation of CLE genes finds 60 thousand newly annotated CLE genes in 2000 analyzed plant genomes. **A)** CLE genes are post-translationally modified into active dodeca peptides, which are bound by LRR-RLKs as a part of signalling in diverse plant developmental processes (some examples are represented in the different bubbles). **B)** Low Conservation Score derived from the EMS2 language model and low expression of CLE genes (red) compared to other gene families. **C)** This study’s pipeline to annotate CLE genes, consisting of a first step with Diamond, followed by gene reconstruction by MAKER2 and filtering. **D)** Percentage of newly annotated (red) or re-annotated *CLE* genes in a subset of species with annotated genomes. Re-annotated *CLE* are further stratified in previously-known (dark blue) and previously-uncharacterized (light blue). Aggregated data across genomes are also summarised and percentage values are shown.

Beyond many past and present efforts to elucidate the functional roles of individual CLE peptides, a long-standing, yet intractable, endeavor has been to study the complex evolutionary histories that have shaped *CLE* family compositions and functions, as this understanding may provide insights into the dynamics and mechanisms driving functional diversification. Highly variable *CLE* family compositions, rapid coding sequence divergence, and generally low expression levels all complicate annotation efforts (Carbonnel, Falquet, and Hazak 2022). In particular, resolving *CLE* family sequence evolution has been hampered by the difficulties associated with aligning short, rapidly evolving proteins, often yielding low-quality alignments that compromise downstream analyses (Goad, Zhu, and Kellogg 2017).

The exponential increase in reference genomes along with alternatives to conventional sequence alignment approaches, has opened opportunities to investigate *CLE* family diversification. As a first step to improve *CLE* annotation, we adopted a method to assess and compare sequence conservation among members and between gene families based on the protein language model EVOLUTIONARY SCALE MODELING 2 (ESM2) (Yeung et al. 2023), bypassing the need for multiple sequence alignment. We first validated that ESM2 was appropriately trained for this task by confirming its ability to detect fundamental elements—such as the dodecapeptide (**Supplementary** Figure 1). We then applied it to the current repertoire of annotated gene families across plant, which revealed that *CLE* genes and other small signaling peptides exhibit much lower conservation and expression compared to other gene families. For example, ribosomal proteins display the highest conservation, whereas *CLEs* are more closely associated with other small signaling peptides (eg. IDAs (Furumizu and Shinohara 2024)) and specific transcription factor families (**Figure 1B**).

This ESM2 analysis validates the extreme diversification of the *CLE* gene family and underscores the annotation challenges inherent to these genes. The more rapid evolutionary dynamics compared to other families and restricted expression profiles of *CLE* genes suggest that the annotation challenges previously observed in models such as Arabidopsis and tomato (*Solanum lycopersicum*) (Carbonnel, Falquet, and Hazak 2022; Carbonnel, Cornelis, and Hazak 2023) are widespread among other plant genomes. In addition, general annotation tools are more likely to miss *CLE* genes due to their short sequences, limited transcriptional support, and low homology scores. To overcome these challenges, we developed a pipeline to comprehensively re-annotate existing and discover undocumented *CLE* family members via a pan-angiosperm scan of over 2,000 genomes spanning over 1,000 species (**Figure 1C**; **Supplementary Table 1**; see **Methods**). We first compiled peptide data from 400 well-annotated species’ genomes to create a dense homology search dataset. Using Diamond, a tBLASTn-like tool (Buchfink, Xie, and Huson 2015), we scanned genomic regions for *CLE*-like sequences. We then directed the annotation algorithm MAKER2 (Holt and Yandell 2011) to those regions, increasing annotation sensitivity by reducing the search space. Finally, we used a Hidden Markov Model to enrich true *CLE* genes by confirming the presence of a signal peptide. This approach identified over 2 million BLAST hits, of which 250,000 were annotated as protein-coding genes by MAKER2, ultimately yielding 60,000 genes classified as *CLEs* (**Figure 1C**).

To evaluate the impact and accuracy of our pipeline, we first applied it to a subset of well-annotated genomes (see https://conservatorycns.com for details). We found that over 40% of these species harbored previously unannotated *CLE* genes, and nearly all species contained mis-annotated *CLE* genes (i.e., genes annotated as protein-coding but not recognized as *CLE* family members) (**Figure 1D** and **Supplementary Table 2**). In well-studied genomes such as Arabidopsis all *CLE* genes were correctly annotated, whereas the close relative *Arabis alpina* had 14 family members that were not annotated, increasing the total number of *CLE* genes in this species to 34.

### Phylogenetic reconstruction of the *CLE* gene family and modeling co-evolution with LRR receptors

In addition to annotation challenges, the short sequences and extreme sequence diversification of *CLE* genes impede the construction of reliable multiple sequence alignments to build robust phylogenetic trees within and across species. One strategy that overcomes this limitation is graphical representations of gene clusters based on reciprocal BLASTp networks (Goad, Zhu, and Kellogg 2017). We attempted to use this approach (Goad, Zhu, and Kellogg 2017); however, standard reciprocal BLASTp algorithms, such as CLANS (Frickey and Lupas 2004), typically capture only closely related relationships and are sensitive to sample size. To address unravel the intricate genetic structure within the pan-angiosperm *CLE* family, we employed *Node2Vec*, a graph-representation embedding method that enabled efficient analysis while minimizing redundancy and noise by taking a random-walk approach (Grover and Leskovec 2016). The resulting high-dimensional embedding was then projected onto a two-dimensional map using PHATE (see **Methods**), thereby preserving both global and local data structures and overcoming the limitations of methods that capture only local relationships (Moon et al. 2019).

The resulting projection revealed distinct clustering patterns that reflect known relationships and give important context to newly annotated genes. For example, CLV3 and close paralogs such as SlCLE9 in tomato and AtCLE40 in Arabidopsis cluster together, demonstrating their sequence similarity (**Figure 2A** and **Supplementary Table 3**). Moreover, hierarchical clustering (see **Methods**) of the complete set of unprocessed CLE protein sequences produced a dendrogram that exhibits major splits associated with amino acid changes in the dodecapeptide (**Figure 2B**). These splits reflect divergent evolutionary trajectories and underscore the central contribution of dodecapeptide sequences in resolving complex phylogenetic relationships within the CLE family. In contrast, other sequence-specific features, such as the composition of the Golgi N-terminal signal sequence (**Supplementary** Figure 2), showed a weaker association, reinforcing that the dodecapeptide is the dominant element shaping the observed evolutionary patterns, owing to its deeply conserved functional role in signalling through cell-surface LRR receptors.

**Figure 2.**
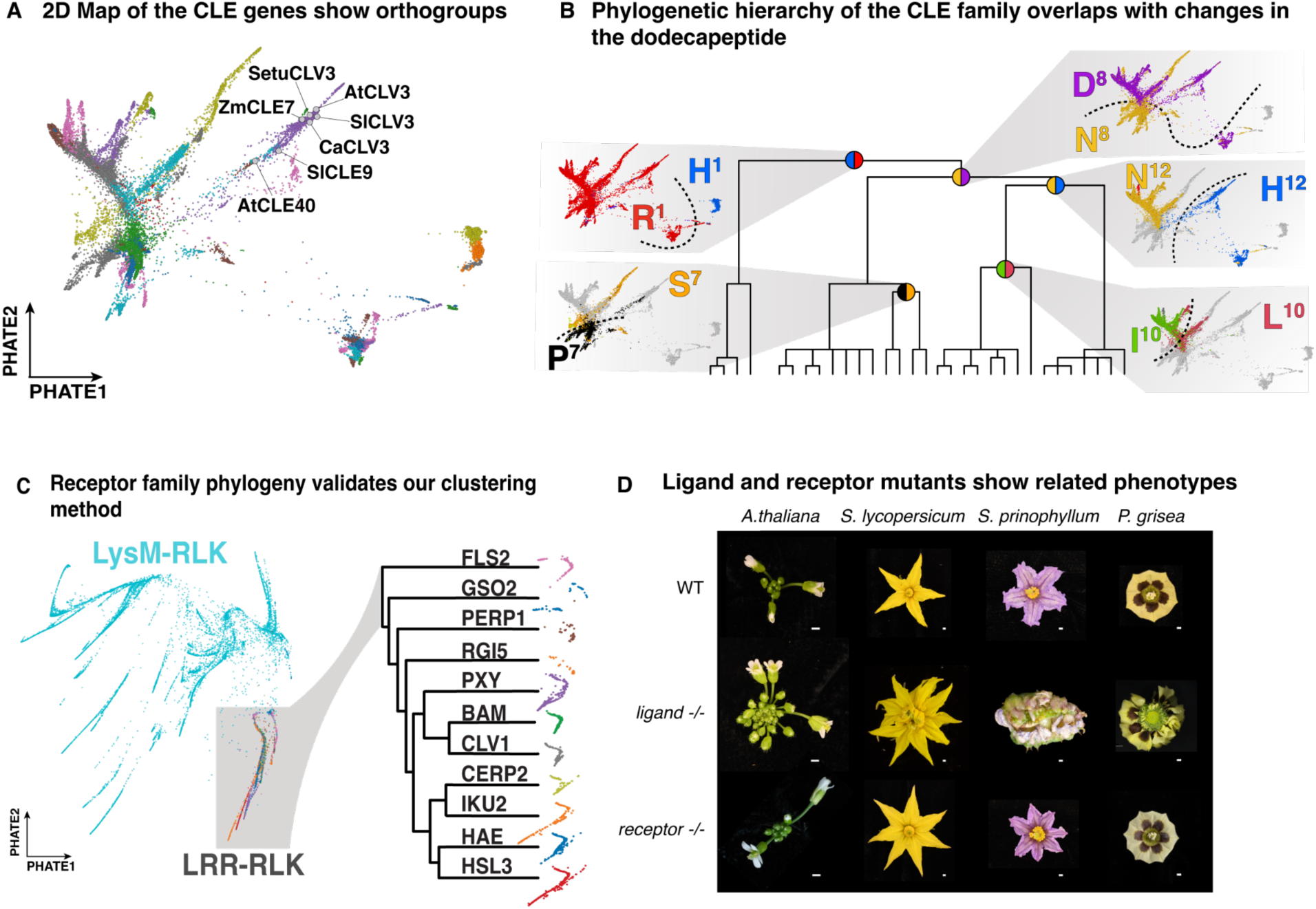
Our pipeline enables new analysis of the evolutionary dynamics of CLE genes. **A)** PHATE Map of the BLASTp pairwise comparison network of *CLE* genes showing different clusters, highlighting known *CLV3* orthologous and paralogous genes that are proximal to each other. **B)** Dendrogram of hierarchical clustering of *CLE* genes showing evolutionary splits and nucleotide substitutions in the functional Dodecapeptide driving them. For each major split, the PHATE map projections of specific amino acid changes at specific positions follow the branching pattern of the dendrogram (as shown based on the orientation of the colors in the circles located at each split point). **C)** PHATE plot of BLASTp pairwise comparison network of LysM-RLK family genes, highlighting LRR-RLK clade and the correspondence of our clustering with previously published phylogenetic relationships. **D)** Representative apical-meristem-derived floral development phenotypes of CLE ligand and receptor mutants in several species, together with wild-type (WT). Ligand mutant corresponding to *Atclv3* in *A. thaliana*, *Slclv3 Slcle9* in *S. lycopersicum*, *Spriclv3a Spriclv3b* in *S. prinophyllum*, and *Pgclv3 Pgcle9* in *P. grisea*. The receptor mutant corresponds to *clv1* mutant in all species. Scale bar is 1 mm.

Given the outcome from applying graph-representation embedding to phylogenetics, we next generated a vector representation of the *LEUCINE-RICH REPEAT RECEPTOR-LIKE* (*LRR-RLK*) gene family, which includes canonical *CLE* peptide receptors such as *CLAVATA1* (*CLV1*) and *BARELY ANY MERISTEM* (*BAM*) (Ogawa et al. 2008; Rodriguez-Leal et al. 2019; Seo et al. 2024). Leveraging the more reliable annotation status of this gene family, we searched for *LRR-RLK* genes in the same subset of 400 well-annotated genomes (see **Methods**). Cluster analysis revealed clear groupings consistent with previous studies (Man et al. 2023), with the embedding precisely distinguishing known clades such as *CLV1*, *BAMs*, and *PHLOEM INTERCALATED WITH XYLEM* (*PXY*) (**Figure 2C**).

The functional relationship and molecular modes of action between CLE peptides and LRR-RLKs are well established, including detailed structural and biochemical analyses (Zhang et al, 2016). Furthermore, comparative genetic studies across species have repeatedly demonstrated that interactions between members of these families are co-evolutionarily stable (Je et al. 2018; Kwon, et al. 2022; Rodriguez-Leal et al. 2019; Seo et al. 2024; Ogawa et al. 2008). For instance, the functions of orthologs of *CLV3* and its primary receptor *CLV1*—whose mutations cause stem cell overproliferation, increased meristem size, and floral organ overproliferation (fasciation)—are deeply conserved, spanning maize, Arabidopsis, tomato and the Solanaceae species *Physalis grisea* (groundcherry) (Ogawa et al. 2008; Rodriguez-Leal et al. 2019; Seo et al. 2024). We also leveraged genome editing in the our recently established *Solanaceae* genetic system, forest nightshade (*S. prinophyllum*) (Benoit et al. 2025) to mutate its *CLV1* ortholog, which caused moderate fasciation like in tomato (**Figure 2D** and **Supplementary Table 4**).

Given this deeply conserved functional relationship, we sought to test whether the graph-based approach can be applied to examine coevolution between receptors and their peptide ligands. A recent study on the interaction between the signalling peptides SCOOPs and their receptor MIK2 showed the power of generative modeling in dissecting interaction mechanisms (Snoeck et al. 2024). Following this methodology, we applied modelling based on Alphafold Multimer to delineate the interaction landscape within the LRR binding region (**Supplementary** Figure 3A). Among putative interacting positions, we observed that certain residues in the receptor domain showed a distribution of amino acid identity per position in the embedding space with clear splits, as observed for the dodecapeptide (see **Methods**). Given the inability to use paired multiple sequence alignments between ligands and receptors to detect co-variation, these graph-embedding-based observations suggest that these CLE peptide and LRR binding domain residues may have been involved in evolutionarily important ligand interactions. A striking example of this relationship is positions 152 and 177, both having asparagine (N) in the CLV1 and BAM clades, but Serine (S) in PXYs (**Supplementary** Figure 3B). In a crystal structure, these residues are predicted to be important for interaction with position 1 of the CLE dodecapeptide, representing a restricted amino acid whose shift in residues mirrors the R–H dichotomy observed at position 1 in our dodecapeptide clustering (H. Zhang et al. 2016). In addition to validating the utility of a graph-representation embedding approach in comparative phylogenetics, this analysis provided insights into possible coevolution of CLE peptides and their receptors.

### Mutational effect analysis of CLE dodecapeptides

Beyond assembling a comprehensive pan-angiosperm collection of *CLE* genes for phylogenetic reconstruction, we asked whether uncovering the full breadth of the family’s sequence diversity could yield additional functional insights by applying our dataset to an emerging area of computational genetics focused on identifying mutational effect (ME) signals embedded within natural variation. Specifically, we applied a Potts model to learn the co-evolutionary patterns among residues within the peptide and predict the effects of mutations from deep sampling of functional sequence data (Riesselman, Ingraham, and Marks 2018; Hopf et al. 2017).

Using forest nightshade *CLV3* (*SpriCLV3*) as a case study, we observed that its mutational landscape exhibits a non-uniform distribution of effects along the 12 amino acids of functional dodecapeptide (**Figure 3A**). Notably, positions 1 and 5(corresponding to glycine and alanine, respectively) demonstrate more neutral mutational effects, suggesting a lower sensitivity to substitutions, consistent with findings in Arabidopsis (Kondo et al. 2006; 2008; Ogawa et al. 2008). To further validate this observation, we benchmarked our tomato SlCLV3-SlCLV1 ligand- receptor model against SSIPe, a hybrid model that combines sequence and structure profiling with force fields (Huang et al. 2020), as well as against docking estimates generated by AlphaFold2, which have proven effective in predicting the impact of amino acid substitutions on docking (Yang, Milas, and White 2022). Initially, our analysis showed that the predicted mutational effects correlated with evolutionary signals derived from BLOSUM matrices (**Supplementary** Figure 4A; R² = 0.026, MSE = 5.43, ρ = 0.4). After regressing out this signal, we observed strong correlations with both the biochemical properties of the substitutions (Sneath Index) (**Figure 3B**; R² = 0.54, MSE = 0.75, ρ = –0.086) and with alterations in binding energy (**Supplementary** Figure 4B; R² = 0.88, MSE = 0.08, ρ = 0.90). Additionally, docking between SpriCLV3 and SpriCLV1 exhibited a moderate correlation with our ME estimates (R² = 0.57, MSE = 0.82, ρ = 0.48) (**Supplementary** Figure 4C).

**Figure 3.**
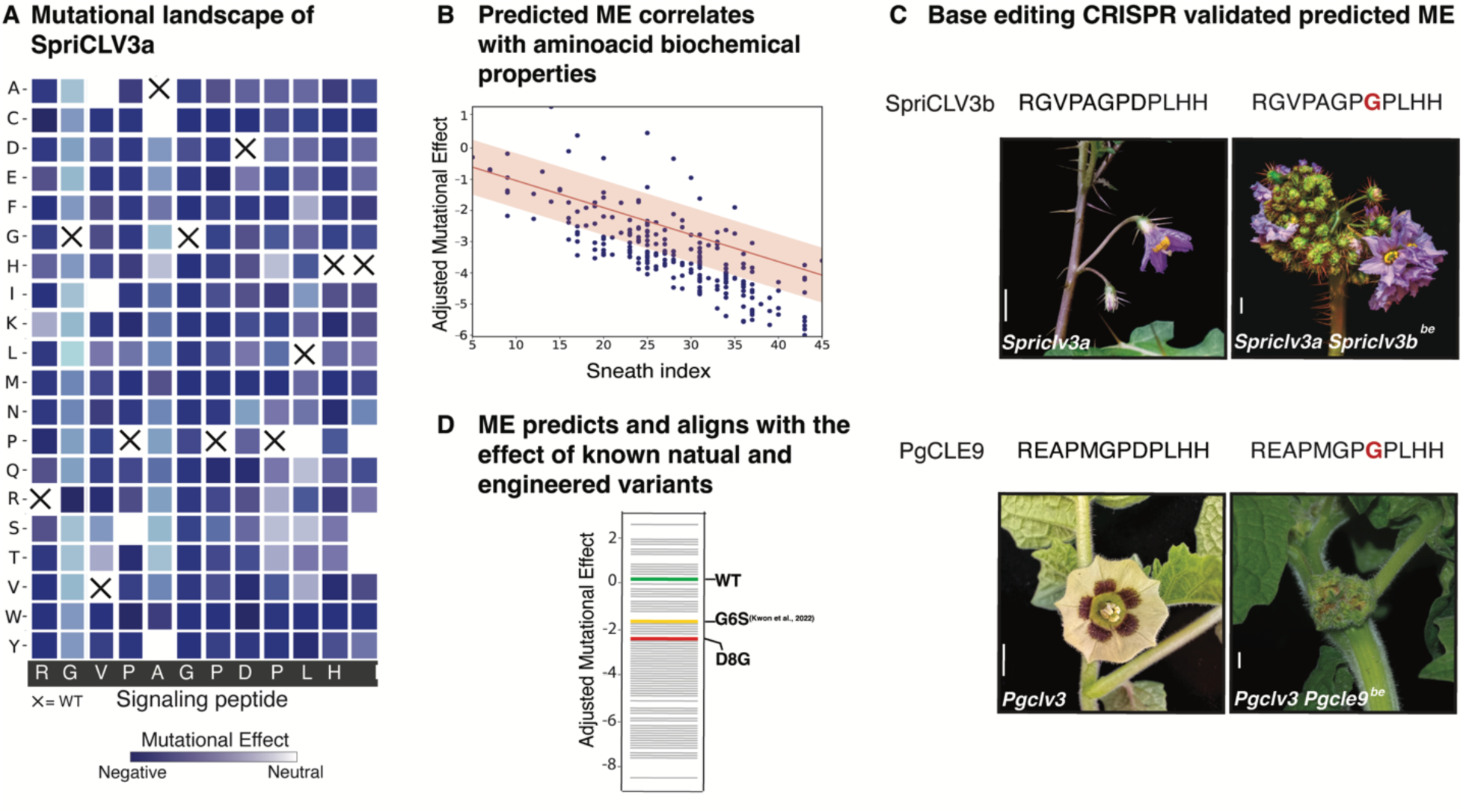
Mutational Effect analysis of the newly expanded *CLE* family allows phenotype prediction. **A)** Mutational landscape of *SpriCLV3* derived from the Pott’s model. Crosses show WT aminoacid position. **B)** Predicted Mutational Effect correlates with Sneath’s index, representing the dissimilarity of the biochemical properties of the amino acids. R² = 0.54, MSE = 0.75, ρ = –0.086 **C)** Dodecapeptide sequence and phenotypes of *CLV3*-clade double mutant base-edited (D8G) plants showing fasciation in two species (*S. prinophyllum* and *P. grisea*) compared to single *clv3* mutants. Base edited residue highlighted in red. **D)** Perceived phenotype strength (WT - green; Moderate - yellow; Severe - Red) of naturally occurring (G6S) and designed substitutions (D8G) correlates with predicted Mutational Effect.

Following exhaustive *in silico* validation, we assessed functional predictions of our approach *in vivo*. Our previous work demonstrated that a naturally occurring amino acid substitution in SlCLE9 (the partially redundant paralog of SlCLV3) from glycine to serine at position 6 weakens peptide function, leading to only partial compensation for loss-of-function mutations in *SlCLV3 (Kwon et al. 2022; Aguirre et al. 2023)*. In contrast, in other *Solanaceae*, such as groundcherry and petunia (*Petunia hybrida*), this residue is maintained as the ancestral glycine, and *CLE9* orthologs are more potent compensators when *CLV3* orthologs are mutated (Kwon et al. 2022). Notably, our modeling similarly predicted a deleterious effect associated with the serine substitution, corroborating our previous genetic findings (**Figure 3C**).

To further test functional predictions from our model, we examined species with differing *CLV3* paralog diversifications. The groundcherry *SlCLE9* ortholog (*PgCLE9*) retains near-complete redundancy with *PgCLV3*, whereas in forest nightshade, *SpriCLE9* was lost but redundancy was restored via a local duplication of *SpriCLV3*. We performed CRISPR base-editing of *PgCLE9* in groundcherry and of the derived *SpriCLV3b* paralog in forest nightshade within their respective *clv3* mutant backgrounds (**Figure 3C**) (Kwon et al. 2022, Benoit et al. 2025). Substitution of glycine with serine at position 8 in both species’ *CLV3* paralogs resulted in a severe fasciation phenotype, supporting another model prediction (**Figure 3D**) and indicating that the amino acid change in this engineered allele exerts a stronger mutant phenotypic effect than the previously characterized hypomorphic G6S change in tomato (**Figure 3D**).

With these analyses, we have demonstrated the predictive power of functional variants through molecular evolution modeling, aligning both computational and empirical evidence. Our approaches and findings highlight the power of extensive meta-analysis in capturing diversity beyond primary model systems and reveal biological insights that can drive deeper dissection of sequence patterns and their functional consequences.

### Assessing mutational burden in predicted *CLE* paralog redundancy detects asymmetric divergence

A current challenge in interrogating the functions of complex gene families via genome editing is the poor predictability of genotype–phenotype relationships among paralogs (Benoit et al. 2025; Iohannes and Jackson 2023). Building on the computational and experimental validation of our ME predictive model, we repurposed it to assess mutational burdens among *CLE* paralogs (see **Methods** and **Supplementary** Figure 5A). Paralog evolution typically starts from complete redundancy, which relaxes selective pressures and permits the accumulation of mutations, eventually leading to paralog divergence in sequence and function. This divergence can occur at both the coding and regulatory levels. Our model’s ability to quantify mutational effects can offer new insights into the impact of protein-level variation. Given the difficulties in defining the products of gene duplication that retain functional relationships, we classified genes as functional paralogs based on both coding and promoter sequence conservation, where most cis-regulatory function is often found (see **Methods**). Using the paralog groups thus identified, we applied our Potts model to quantify specific paralogs that accumulated more deleterious mutations.

In a broader analysis, we compared the mutational burden of paralogous dodecapeptides versus family-wide homologous dodecapeptides, while also accounting for duplication age based on synonymous substitutions. Our results indicate that closely related, derived paralogs of ancestral family members tend to accumulate more deleterious substitutions than non-paralogous genes **Supplementary** Figure 5B; see **Methods**). These findings are consistent with the theoretical expectation that gene duplication and redundancy allows greater mutation accumulation via relaxed selection (Wagner 2008).

Taking into consideration both coding and promoter sequences, we can more accurately place *CLE* genes within their proper phylogenetic context, thereby enabling improved prediction and dissection of paralog relationships and divergence. Tomato *SlCLV3* and *SlCLE9* exemplify a scenario where their divergence, mediated by both amino acid substitutions and promoter degradation, has led to functional drift while preserving compensatory interactions (Kwon et al. 2022). To further explore these patterns, we focused on another pair of tomato *CLE* paralogs, *SlCLE7* and *SlCLE24*, orthologous to *AtCLE45 and AtCLE33* involved in vasculature formation (Carbonnel, Cornelis, and Hazak 2023). Analysis of the *SlCLE7-SlCLE24* clade (comprising only two members) revealed an analogous paralog relationship to that of *SlCLV3–SlCLE9*. Despite not showing the accumulation of deleterious coding mutations between the pair, with three changes having a neutral net effect, these paralogs showed asymmetrical promoter degradation, with the *SlCLE7* promoter showing less conservation of the region shared between the two paralogs once compared among its orthologs relative to the promoter conservation among the *SlCLE24* orthogroup (**Figure 4A**, see **Methods**). In line with this observation, *SlCLE7* and *SlCLE24* share nearly identical expression patterns across tissues, but *SlCLE24* exhibits higher expression (**Figure 4B**), suggesting *SlCLE24* paralog dominance in this predicted unequal redundancy relationship(Benoit et al. 2025).

**Figure 4.**
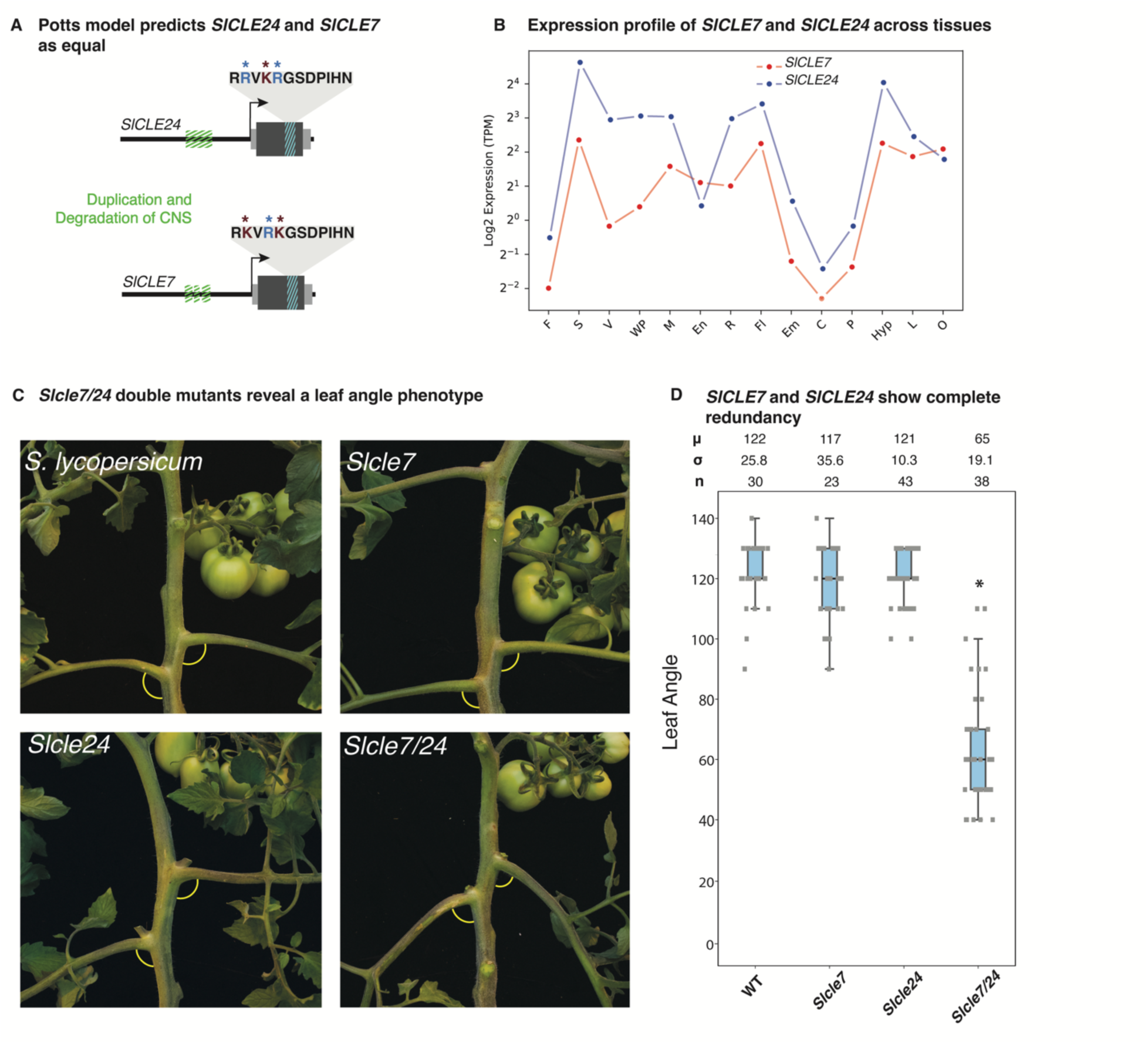
*CLE7* and *CLE24* redundantly control petiole angle in tomato. **A)** Scheme of the variation present between *SlCLE7* and *SlCLE24*, showing sequence variation in both the CDS and the promoter region of both genes. Substitutions in the dodecapeptide with a predicted negative effect are marked in red, while positive changes are in blue. Cumulatively, the net difference between *SlCLE7* and *SlCLE24* is close to zero. **B)** Expression (Log2 transcripts per million, TPM) of *SlCLE7* and *SlCLE24* in different tissues (F: Fruit, S:Stem, C:Vasculature, WP:Whole Plant; M: Meristem, EN: Endosperm, R:Roots, Fl:Flowers, Em:Embryo, C:Calli, P:Pollen, Hyp:Hypocotyl, L:Leaves, O:Ovary. Red, *SlCLE7*; Blue, *SlCLE24*. **C)** Representative petiole angle phenotypes of wild type tomato, *Slcle7* and *Slcle24* single mutants and *Slcle7 Slcle24* double mutants. **D)** Quantification of petiole angle phenotype in wild type tomato, *Slcle7* and *Slcle24* single mutants and *Slcle7 Slcle24* double mutants. ‘μ’ represents the mean, ‘σ’ refers to standard deviation and ‘n’ indicates the sample size used. ‘*’ indicates a statistical difference < 0.05.

To better understand the relevance of regulatory and coding sequence conservation in the predicted redundancy relationship of *SlCLE24* and *SlCLE7*, we simultaneously mutated both genes using CRISPR/Cas9 gene editing. A screen of progeny from first-generation transgenic (T0) plants revealed a conspicuous change in leaf angle. While wild-type leaves exhibited an angle of 110– 130° relative to the main shoot, plants with null mutations in both *Slcle7* and *Slcle24* (double mutants) showed a substantially reduced leaf angle of 60–90° (p < 0.001) (**Figures 4C–D**).

Notably, this reduction of leaf angle mirrors that of the previously characterized mutant *fasciated and branched 2* (*fab2*) mutant (Jeong et al. 2024), defective in an enzyme involved in arabinosylation of the tomato CLV3 dodecapeptide and of other CLEs in Arabidopsis (**Supplementary** Figure 6), suggesting a role for FAB2-mediated modification of *SlCLE24- SlCLE7* in this developmental syndrome (Jeong et al. 2024). This phenotype is absent in single mutants of each gene, indicating that the regulatory divergence reflected by differences in promoter sequences and gene expression is insufficient to compromise functionality; rather, the inherent strength of the peptides, as indicated by the lack of deleterious mutations, preserves their compensatory roles. In contrast to the situation observed for *SlCLV3* and *SlCLE9*, where promoter degradation significantly alters their interaction, the functional redundancy between *SlCLE7* and *SlCLE24* remains balanced, despite the observed regulatory sequence degradation and signal of paralog dominance from our sequence analysis.

### Dissection of the complex genetic interactions in two additional CLE clades

While *SlCLE7-SlCLE24* provides a relatively simple paralog pair to investigate how sequence divergence translates into functional drift, many other *CLE* family clades in tomato display considerably greater genetic complexity. For example, two larger *CLE* clades identified through our computational analyses, designated R1D8H12 and R1N8N12 based on amino acid composition and distribution on dendrogram (**Figure 5B**, **Supplementary Table 3**), offer opportunities to dissect genetic architectures and patterns of paralog redundancy in greater depth. Assuming higher-order redundancy among members of these clades, we developed a combined forward and reverse genetics approach using multiplex CRISPR/Cas9 editing that targets all members of a given clade. We generated multiple independent transgenic mutant populations and performed bulk short-read sequencing approach to genotype both segregating and fixed lines, establishing a method for associating engineered collections of mutations with emergent phenotypes (see **Methods** and **Figure 5A, 5C**). This strategy addresses the limitations of simpler gene editing approaches, which often fail to resolve the intricate redundancy patterns among numerous family members.

**Figure 5.**
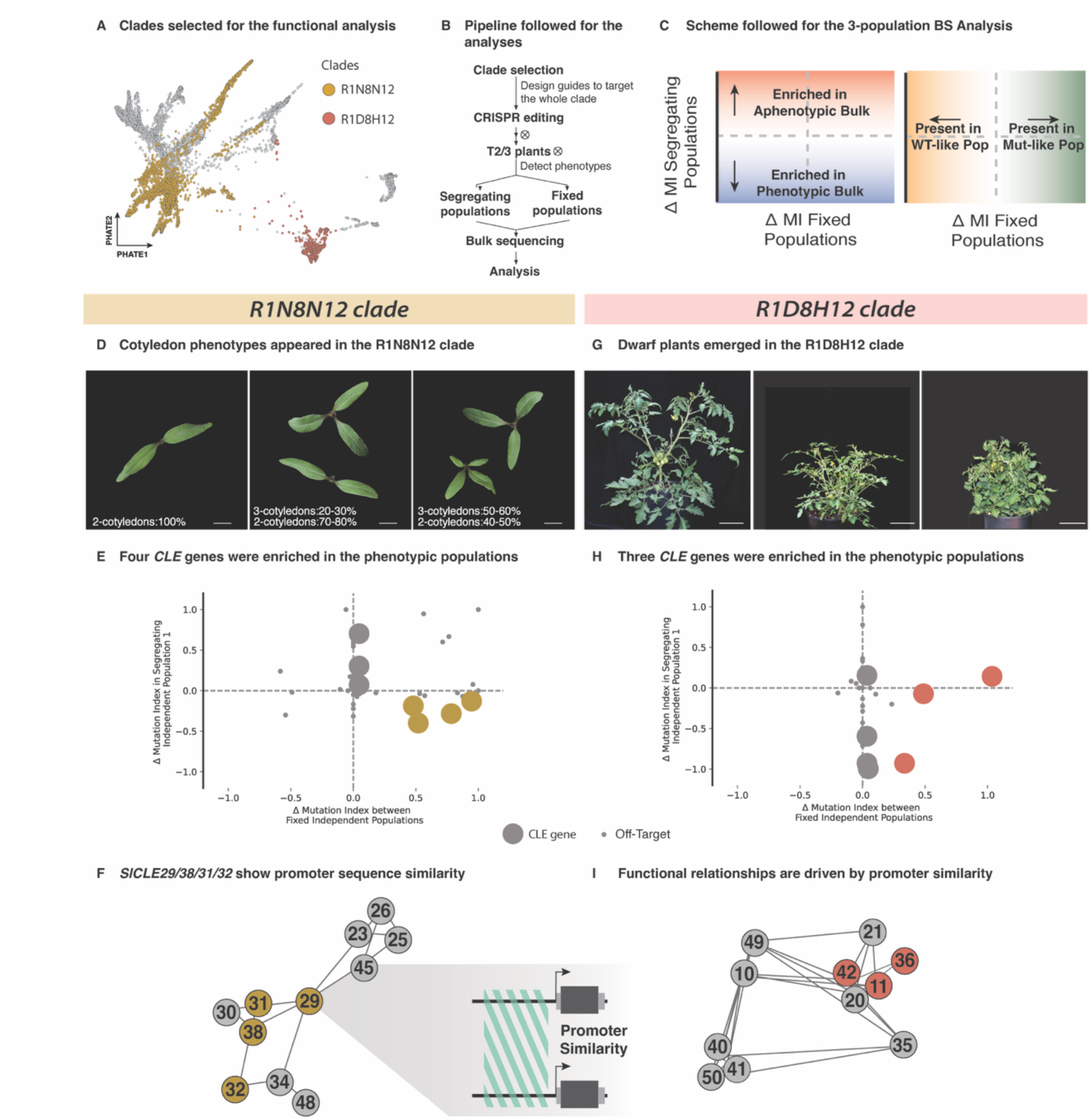
Promoter variation correlates with CLE family functional relationships shaping phenotypes in complex clades. **A)** Pipeline to study gene relationships in complex clades, starting with CRISPR-Cas9 gene editing of the whole clade followed by phenotyping, generation of segregating and fixed populations, and bulk sequencing. **B)** PHATE plot of *CLE* genes, highlighting the two clades selected for analysis, R1N8N12 in dark yellow and in R1D8H12 salmon red. **C)** Scheme of the output of the 3 population bulk sequencing approach used to analyze candidate genes. Presence in the bottom right section indicates enrichment in the mutant phenotypic classes. Mutational index, MI **D, G)** Representative phenotypes obtained by whole-clade gene editing of the two studied clades. R1N8N12 mutant plants revealed cotyledon defects with abnormal presence of extra embryonic leaves. R1D8H12 mutants revealed a more petite, compacted and altered development compared to the aphenoticic populations. **E, H)** Analysis of candidate genes from the bulk sequencing data, showing in the bottom left quarter the genes associated with the observed phenotypes. Small dots refer to offtargets while bigger dots are CLE genes **F, I)** Promoter relationship network, highlighting in color the candidate genes obtained by the analysis. Each node in a promoter of a specific *CLE* gene and each edge is the similarity value between those promoters.

As shown in **Figure 5D**, the R1N8N12 mutational pool was characterized by a multi-cotyledon phenotype with variable expressivity. In one population, 30% of individuals exhibited three cotyledons, whereas in another group, 50% displayed this trait, with occasional instances of four cotyledons. Notably, the Arabidopsis gene *AtCLE19*—a member of this superclade—has been reported to be cotyledon-specific, with mutations causing cotyledon defects (Xu et al. 2015). To determine whether the tomato ortholog or other clade members contribute to this phenotype, we applied our bulk sequencing approach and identified four *CLE* genes as enriched candidates based on their combined mutational profiles (*SlCLE29*, *SlCLE31*, *SlCLE32*, and *SlCLE38*) (**Figure 5E**, and **Supplementary Table 5, Supplementary Table 7** and **Supplementary Table 7** for genotyping information). Importantly, analysis of our integrated datasets, encompassing mutation distances and non-coding sequence information, revealed that promoter similarity predominantly explains the high-order mutational patterns and associated phenotypes observed in the CRISPR lines. Indeed, the top candidate genes clustered tightly within the promoter similarity network, showing a statistically significant pattern that would not occur by chance (**Figure 5F**).

A similar pattern was observed for the R1D8H12 clade, where plants from different populations exhibited vegetative defects characterized by thinner stems and less complex plant architecture with variable severity (**Figure 5G**). Bulk whole-genome Illumina sequencing of these populations revealed enrichment of *SlCLE11*, *SlCLE36*, and *SlCLE42* (**Figure 5H**, and **Supplementary Table 9, Supplementary Table 6** and **Supplementary Table 7** for genotyping information). Intriguingly, like the putative causal mutations from the R1N8N12 clade, these three genes clustered closely in the promoter similarity network rather than by mutational effect variation and expression that on the other hand show contradicting patterns, including for *SlCLE11* that shows the strongest expression profile and the peptide with the highest amount of deleterious mutations (**Figure 5I** and **Supplementary** Figure 7A and B).

## Summary

In this study, we employed an interdisciplinary approach, from custom *de novo* gene-annotation pipeline to CRISPR mutagenesis, to decode the diversification of CLE signaling peptides across flowering plants. This strategy not only yielded broad insights into *CLE* family evolution, but also established a framework for studying other rapidly evolving gene families and for dissecting the co-evolutionary dynamics including between their interacting partners, for example, the reciprocal changes between CLE peptides and their LRR receptors. By bridging the gap between model and non-model species our deep pan-genomic sampling empowered by over 2,000 angiosperm genomes allowed us to disentangle the relative contributions of changes in coding versus cis-regulatory sequences in the diversification of paralogs and their redundancies. By juxtaposing *SlCLE7/24*, whose nearly invariant coding sequences underpin a classical redundancy, with the R1N8N12 and R1D8H12 clades, in which highly conserved promoters rather than protein identity sustain overlapping functions, we expose two complementary routes by which paralogs buffer development; these contrasting cases mirror a broader pattern in which coding fidelity often supports robust, dosage-sensitive compensation within recent duplicates, whereas promoter conservation enables older, more coding sequence-diverged paralogs to retain shared context-specific expression patterns and functions (Wang et al. 2021; Wong et al. 2020; Kvon et al. 2016; Galupa et al. 2023).

Our study demonstrates that a large number of genomes spanning a broad evolutionary space can improve predictions on gene divergence happening over short timescales. Indeed, through comparative functional analyses of the *CLV3* clade in *Solanaceae* (tomato, forest nightshade, and groundcherry), representing less than 50 million years of evolution (J. He et al. 2023), we found that phenotypes derived from CRISPR-mediated base editing of the dodecapeptide aligned with predicted mutational effect modelled based on the full sequence diversity of this gene family. This demonstrated that global modelling of gene family sequence diversity can predict local patterns of its genotype-phenotype landscape among cohorts of paralogs within those gene families. The phenotypic effect of mutations is the mechanism by which gene families expand and fine-tune their functional repertoire while preserving essential roles. By integrating deep comparative genomics with predictive mutational modeling and targeted editing we provide a roadmap for forecasting when redundancy will fail, how compensation evolves, and how hidden paralog variation can be leveraged to reshape plant form and function.

## Materials and Methods

### CLE and LRR Discovery and de novo annotation

All previously annotated CLE genes were searched in available resources of the Conservatory project, which collected the genomes of more than 300 species, assessed to have a complete annotation. This dataset was then used as a query to search using Diamond (Buchfink, Xie, and Huson, 2015) in all the available plant genomes on NCBI with an assembly status of Scaffold or Chromosome [Diamond parameters: –ultra-sensitive –masking 0 –iterate]. The identified hits were isolated, including 1000 bp upstream and downstream (customized Julia script available on Github). These sequences were subsequently analyzed by MAKER2 (Holt and Yandell 2011) with dynamic parametrization based on the genome of origin. The predicted proteins were then assessed for being true CLE genes by checking for the presence of a signaling peptide and the CLE motif using InterProScan and HMMER [hmmsearch with parameters: hmmsearch –max -T 0] (Jones et al. 2014; Finn, Clements, and Eddy 2011). The newly discovered CLEs were then compared with BEDTools (Quinlan and Hall 2010) to the original source of the well-curated annotation files screened by the Conservatory project to evaluate whether the developed pipelined captured previously unknown genes.

For LRR genes, we run HHMER [hmmsearch with parameters: hmmsearch –max -T 0] for all the proteins from the conservatory genomes. We then filtered for those proteins with > 10 LRRs and a kinase domain.

### Phylogenetic Reconstruction and Conservation Analysis

We generated an all-by-all reciprocal BLASTp comparison using CLANS 2.0 (Frickey and Lupas 2004, available on CLANS2.0 https://github.com/inbalpaz/CLANS). We constructed an undirected network using − log10(BLASTp E-value) as edge weight. We then pruned the network so that each node maintained the top 500 connections (customized Python script available on Github). Node2Vec (Grover and Leskovec 2016), implemented in SNAP (Leskovec and Sosic 2016), was then used to vectorize the outputted graph [-l:200 -r:600 -k:100 -e:1 -w]. The vectorized network in 128 dimensions was then projected into a lower-dimensional space using PHATE (Moon et al. 2019). To detect paralogy and homology, we implemented the Leiden algorithm (customized Python script available on Github). We adopted multiple increasing resolution parameters and reconciled all the outputs into a hierarchical structure using the Multi-resolution Reconciled Tree in R (Peng et al. 2021). To assess and resolve the over-clustering at the leaf level in the generated tree, we implemented a RandomForest-based permutation test (similar to the system used in CHOIR (Petersen, Mucke, and Corces 2024) using the dodecapeptide as a 12-dimensional label space.

Sister leaves that failed the permutation test were pruned. To resolve the polytomy at higher nodes, we adopted a parsimonious entropy minimization test based on the dodecapeptide amino acid composition (customized Python script available on Github). To integrate promoter information into the structure to better delineate paralogy and homology, we first constructed a kNN graph based on the vectorized BLASTp network. Each node was unlabeled except for those CLE genes that overlapped with the Conservatory project. We extracted from the Conservatory project the promoter conservation levels with other CLE genes within the same genome or between species. CLE genes with shared promoters were labeled as a joint or the group and labelled accordingly. To sort the remaining CLE genes that did not have promoter information available, we applied a label propagation algorithm to the partially annotated kNN graph (customized Python script available on Github).

For LRR gene classification, we took the sequences from and classified in their specific phylogenetic classes, and built a random-forest classifier for each phylogenetic clade based on pairwise comparison. We then applied this classification model to the discovered sequences in this study and overlapped the distributions of these predicted labels to the outputs of leiden cludstering, observing clear overlaps.

For the conservation analysis via EMS2 we adopted the method in REF, adopting model. The conservation score was calculated as the proportion of the positions with a site-level value above 0.6. All scripts are available at https://github.com/LippmanLab.

### Mutational Effect Estimation and Validation

To assess the mutational burden between paralogs or homologs, we extracted all the CLE motives from our dataset based on the match derived from hmmsearch. The subsequent list of sequences was treated as a gapless multiple sequence alignment and inputed in EVmutation (Hopf et al. 2017) following the author’s guidelines and implemented in pipeline available on Github. The derived results were validated computationally by focusing on SlCLV3. The three-dimensional structure of SpriCLV3 interacting with its LRR receptor SpriCLV1 was obtained using AlphaFold-Multimer (default parameter - relaxed) (AlphaFold-Multimer). The generated structure was cross-validated with known information about the physical interaction between CLE peptides and LRR receptors (Morita et al. 2016). Once the structure was validated, confirming the power of AlphaFold-Multimer to generate outputs resembling the expected interactions, we performed an in silico saturation mutagenesis using SSIPe (Huang et al. 2020) to evaluate binding affinity and AlphaFold-Multimer to assess docking, following the method described in (Yang, Milas, and White 2022). In addition to these measurements, ME values were also compared to the biochemical properties of the substitutions (Sneath Index). The resulting quantifications were used in a regression analysis to evaluate our predicted mutational effects. The code is available at

### Plant materials, Growth Conditions and Phenotyping

As previously described in (Ciren, Zebell, and Lippman 2024), seeds of wild type Solanum lycopersicum (cultivar M82, LA3475), Solanum prinophyllom and Physalis grisea (ZL05) were used. Seeds were directly sown in soil in 96-cell plastic flats and grown to 4-week-old seedlings in the greenhouse.

Seedlings were then transplanted to 4L pots in the greenhouse for crossing and bulking purposes or directly to the fields at Cold Spring Harbor Laboratory, New York. Green-house conditions are long-day (16 h light, 26-28 °C / 8 h dark, 18-20 °C; 40-60% relative humidity) with natural light supplemented with artificial light from high pressure sodium bulbs (250 umol m−2 s−1). Plants in the fields were grown under drip irrigation and standard fertiliser regimes, and were used for quantifications of inflorescence branching, sepal length, and fruit shape. Quantitative phenotypic data were collected manually in fields and greenhouses. Raw leaf angle data are in **Supplementary Table 11**.

### Expression Analysis

Publically available RNA-seq datasets for tomato and other species were used (**Supplementary Table 10**). For tomato, a similar approach used in (Benoit et al. 2025) was implemented: raw reads were realigned to the reference genome and TPM quantified. Only samples with more than 50% uniquely mapped reads were retained for subsequent analysis. Further filtering was applied based on the Spearman correlation between tissue replicates, and removed samples with low correlation (0.75 or below).

### Genome Editing

As previosly described in (Ciren, Zebell, and Lippman 2024), CRISPR-Cas9 mutagenesis and generation of transgenic tomato plants were performed following our standard protocol. Briefly, guide RNAs (gRNAs) (listed in **Supplementary Table 8**) were designed using the Geneious Prime software. For Cas9 multiplex editing, the Golden Gate cloning system was used to assemble the binary vector containing the Cas9 and the specific gRNAs. For Base editing, vectors were constructed through a modular GatewayTM assembly, as described previously (Invitrogen). Final binary vectors were then transformed into the tomato cultivar M82 by Agrobacterium tumefaciens-mediated transformation through tissue culture.

Regenerated plants and First-generation Plants were genotyped in the target regions through primers designed in Geneious Prime software (listed in **Supplementary Table 8).**

### 3-Population Analysis

As described, Second- and Third-generation transgenic plants (T2/3) were genotyped via Illumina-based WGS on NextSeq 2000 P3 sequencing platform (Illumina). Plants showing a phenotype were bulked together and sequenced. Reads were aligned to the M82 Genome (Alonge et al. 2022) via BWA (Li and Durbin 2010). Polymorphisms were called with Snpift (Cingolani et al. 2012) (**Supplementary Table 6** for *CLE* gene specific alleles and **Supplementary Table 7** for the entire Snipft output) and analysed in a customized Python script based on comparison of a segrating population against 2 fixed populations, wt-like and mutant-like.

## Acknowledgments

We thank members of the Lippman laboratory for their support and feedback, and T. Mulligan, K. Schlecht, S. Qiao for assistance with plant care.

## Data Availability

All alignments, trees, rate test results, selection test results, and phenotyping data are available on the Lippman lab’s GitHub

---

**Supplementary Figure 1.**
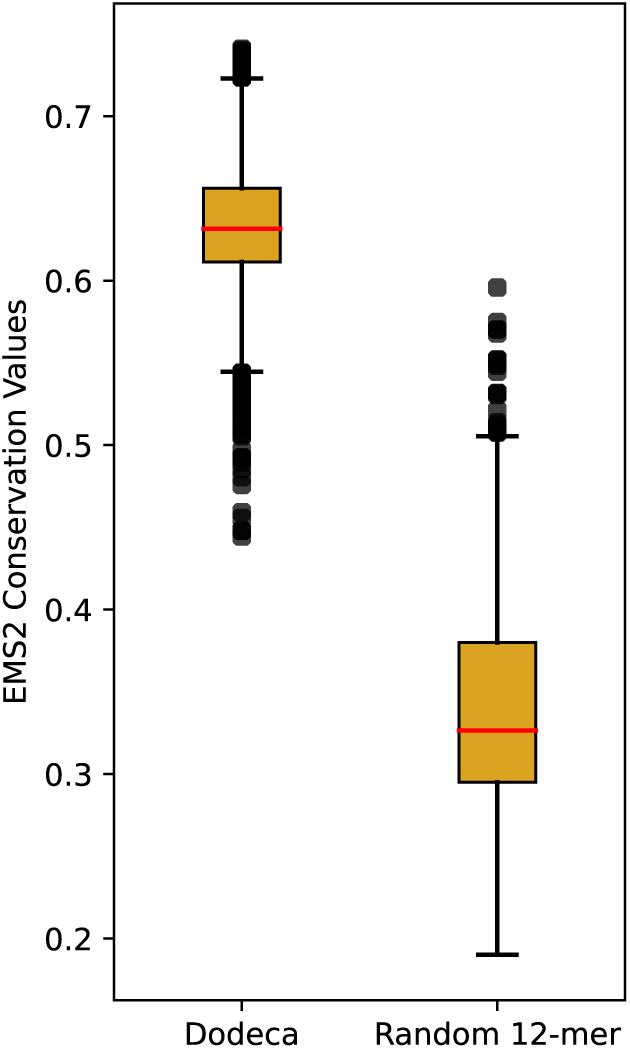
EMS2 Model recognizes CLE Dodecapeptides. The outputted Conservation. The outputted Conservation Value from EMS2 is higher for the section of the CLE protein encoding for the functional 12-mer motif.

**Supplementary Figure 2.**
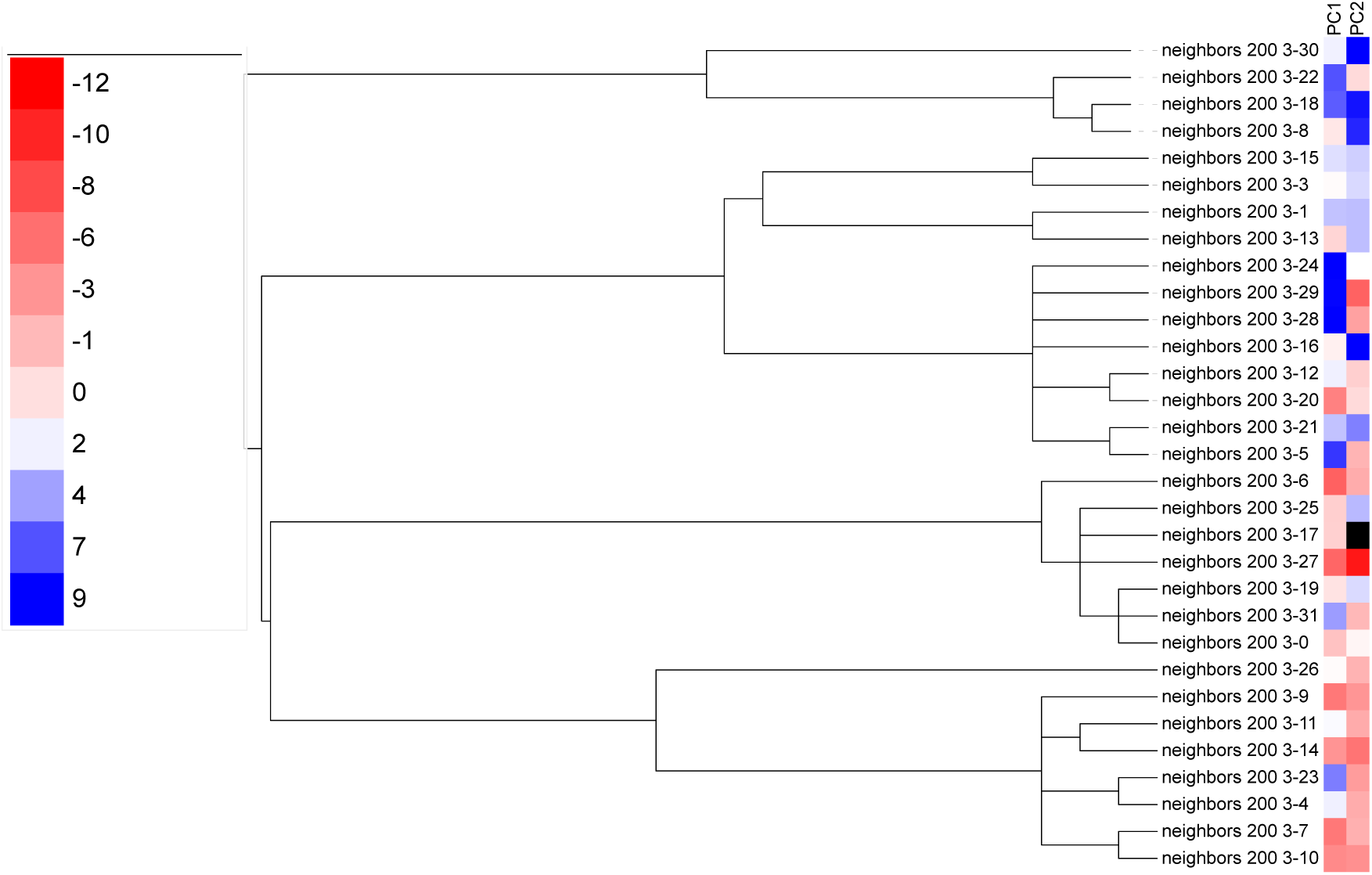
Golgi Signalling peptide sequence Composition in CLE genes. PC1 and PC2 values for Golgi Signalling peptide sequence Composition reveals some patterns given the reconstructed phylogenetic structure.

**Supplementary Figure 3.**
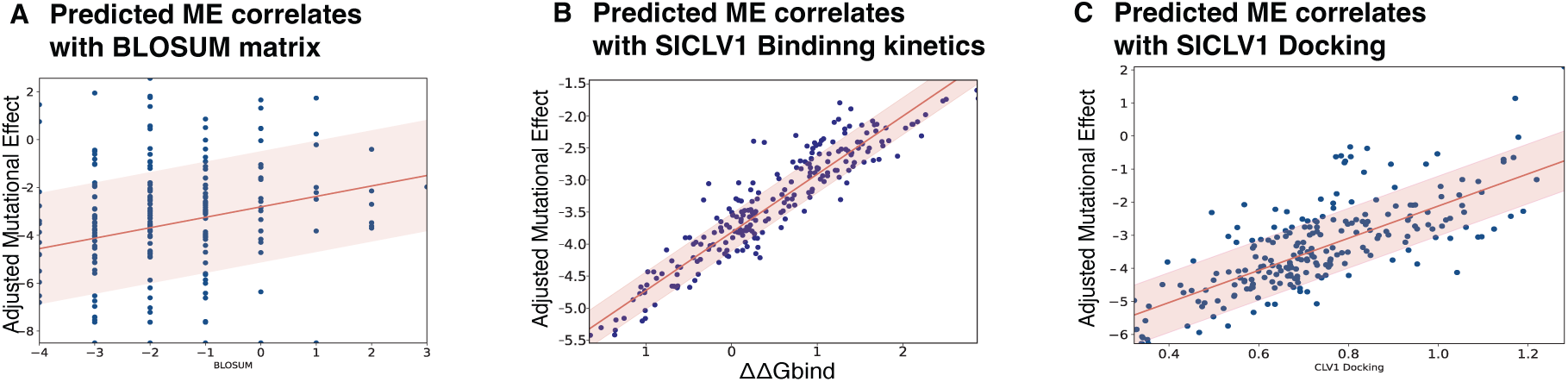
Mutational Effects correlate with relevant biological aspects of CLE biology. A) Mutational Effect prediction correlates with BLOSUM substitution matrix. B) Mutational Effect prediction correlates with Kinetics values derived from SSIPe. C) Mutational Effect prediction correlates with Alphafold Docking scores.

**Supplementary Figure 4.**
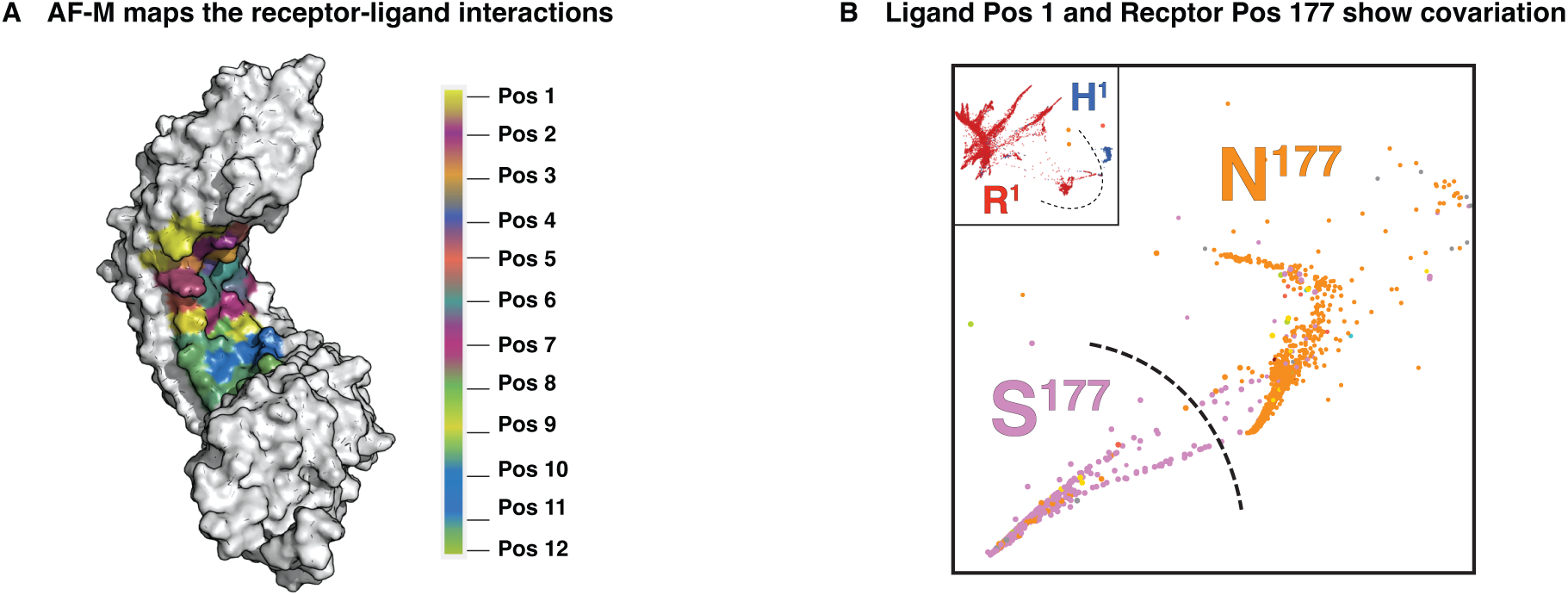
Our pipeline enables new analysis of co-evolutionary dynamics between CLE genes and LRR receptors. A) Alpha-Fold Multimer structure of CLE receptors, highlighting the consensus CLE binding surface per residue derived from aggregated simulations with all the possible CLE dodecapeptides. B) PHATE plot of the CLV1-BAM-PXY receptor genes and CLE genes showing a similar covariation pattern between ligand position 1 (H/R) and receptor position 177 (N/S).

**Supplementary Figure 5.**
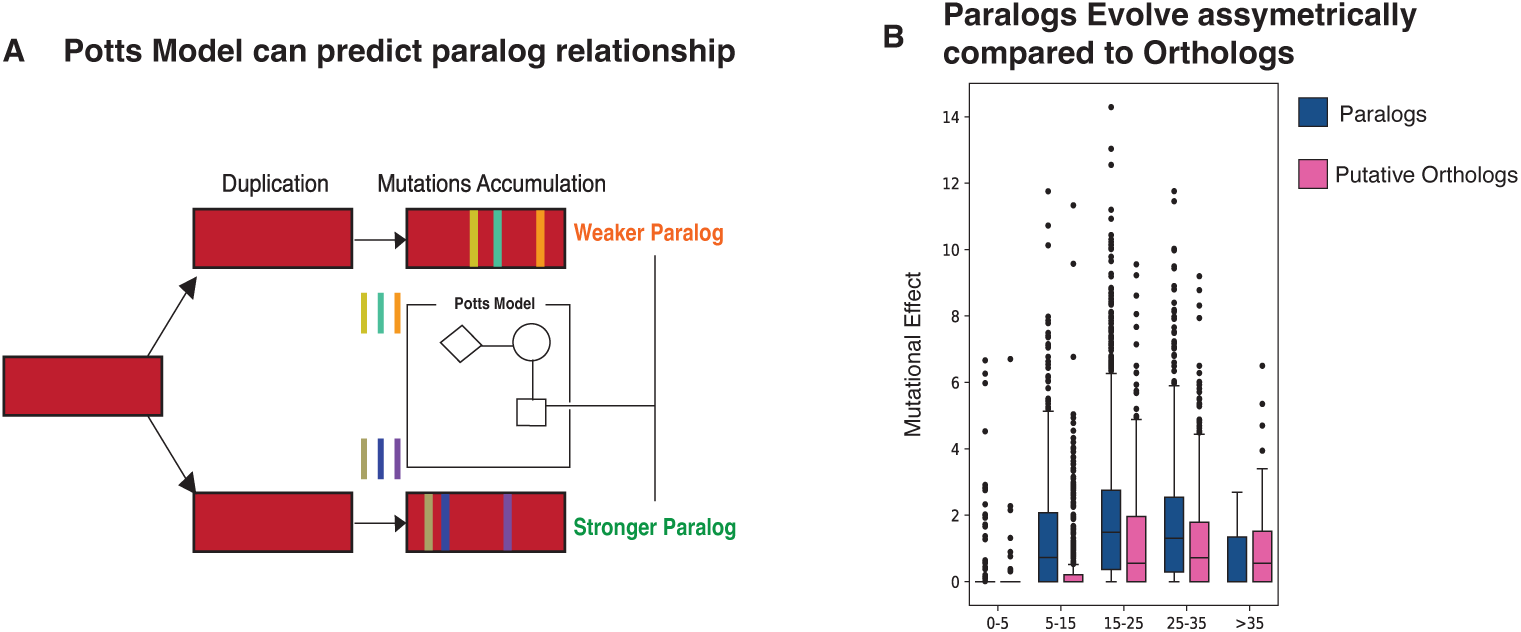
Paralogs have a higher Mutational Burden than Singletons. A) By treating paralogs as the same gene with differentiating mutations, the trained Potts model can interpret these amino acid changes and determine which paralog accumulated the most deleterious amino acid changes. B) Overall, within-species paralog comparisons have a higher mutational burden than between-species singletons.

**Supplementary Figure 6.**
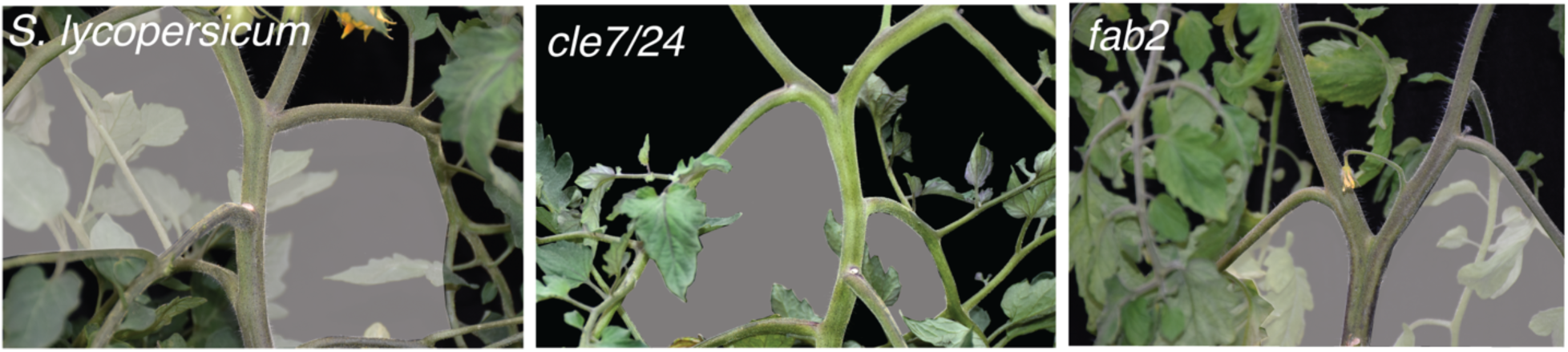
***cle7/24* mutant phenocopies *fab2* mutant, an enzyme involved in the post-translational modification of these peptides.**

**Supplementary Figure 7.**
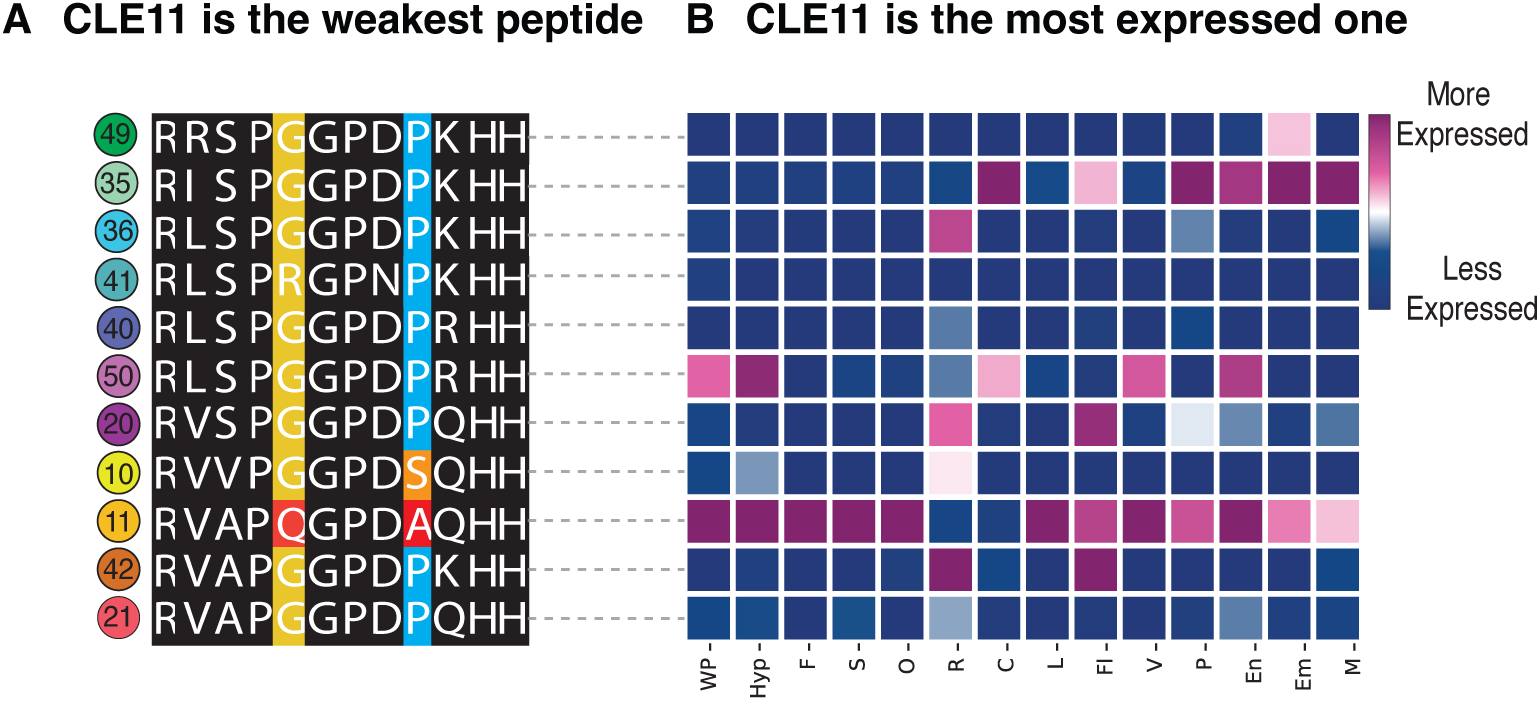
Protein and Expression Analysis of the R1D8H12 Clade. A) Their dodecapeptide composition does not show major variability, except for CLE11 having key amino acid substitutions occurring in two relevant positions. B) Despite being the weakest peptide, CLE11 is the member of this clade showing the highest level of expression, except in roots where CLE42 is the highest expressed gene.

